# Exercise-induced Sweat Promotes Wound Healing in Diabetic Foot Ulcers

**DOI:** 10.64898/2026.04.11.717882

**Authors:** Mengyuan Zhao, Yvxin Tong, Huilin Yao, Jing Cao, Tiantian Liang, Qintong Fei, Maodi Liang, Bojun Yang, Mingjie Sun, Cuizhe Wang, Jun Zhang, Qinghua Cui

**Author notes:** To whom the correspondence should be addressed: Jun Zhang., Cuizhe Wang., Qinghua Cui. These authors contributed equally to this work (Mengyuan Zhao, Yvxin Tong, Huilin Yao, Jing Cao, Tiantian Liang).

## Abstract

Persistent hyperglycemia impairs wound healing in diabetic patients, and severe cases may even lead to disability or death. Glycemic control alone cannot effectively prevent the occurrence of diabetic foot ulcers, a serious complication of diabetes. However, safe, efficient, and cost-effective therapies remain unavailable and are urgently needed. Using a novel sports medicine paradigm, we predicted, based on reverse-transcriptomics, that exercise-induced sweat has the potential to promote would healing in diabetic foot ulcers. Subsequent animal experiments demonstrated that sweat can indeed promote re-epithelialization and collagen deposition, upregulate the expression of the proliferation marker Ki-67, the angiogenesis marker CD31, and α-SMA, and significantly accelerate wound healing in a mouse model of diabetic foot ulcers. This study provides a new direction for sports medicine and offers a novel therapeutic strategy for patients with diabetic foot ulcers.

## Introduction

In 2024, approximately 589 million adults worldwide were living with diabetes. In the same year, more than 3.4 million people died from diabetes or its complications^1^. Due to the long disease course and insidious symptoms of type 2 diabetes (T2DM)^2,3^, about half of adult diabetic patients remain undiagnosed, leaving them in a state of persistent hyperglycemia and significantly increasing their risk of life-threatening and disabling complications^4,5^.

For patients with diabetes, diabetic foot ulcers (DFUs) are one of the most clinically devastating and severe chronic complications. Their core characteristics are characterized by the “four highs and one low” pattern: high incidence, high recurrence rate, high risk of amputation, high mortality, and low cure rate, imposing a heavy burden on patients, families, and the global healthcare system^6-8^. DFUs are caused by multiple pathophysiological factors, including insulin resistance, impaired tissue regeneration, angiopathy, neuropathy, and inflammation^9^. These pathological changes significantly affect various important mechanisms of normal wound healing, potentially leading to impaired wound healing and progression to DFUs^10,11^. Therefore, effectively promoting wound healing and inhibiting its malignant progression is not only a core target for the clinical treatment and drug development of DFUs but also a key breakthrough for reducing the amputation rate and improving patient prognosis. Currently, the most effective strategy for reducing the incidence of diabetes and DFUs is to maintain optimal glycemic control^12^. However, with advancing age, pancreatic β-cell function progressively declines and insulin resistance worsens in diabetic patients, who are frequently complicated by hypertension and hyperlipidemia, making long-term stable glycemic control increasingly challenging^13,14^. Meanwhile, the clinically available wound-healing-promoting agents and therapeutic approaches are not only costly but also associated with substantial limitations^15,16^. Therefore, the development of novel therapeutics with high efficacy, safety, and cost-effectiveness to provide superior treatment options for DFU patients has become a critical scientific issue urgently to be addressed in the field of diabetic complications, which also offers a clear objective and practical significance for the present study.

Previous studies have reported that sweat is not merely a metabolic waste excretion, but a complex biological fluid system rich in a variety of bioactive small proteins and peptides^17^. To date, numerous functional proteins have been identified in sweat, including immunoglobulin A (IgA)^18^, the interleukin family (IL-1, IL-6, IL-8)^19,20^, tumor necrosis factor-α (TNF-α)^21^, and transforming growth factor-β receptor (TGF-βR)^22^, as well as various antimicrobial peptides (AMPs)^23^. These components exert antimicrobial and anti-inflammatory activities and may be closely associated with cutaneous immune defense and tissue repair^24,25^. In this study, using a novel sports medicine paradigm, we predicted, based on reverse-transcriptomics, that exercise-induced sweat has the potential to promote would healing in DFUs. Therefore, as a natural product, the present study hypothesizes that sweat may inhibit the progression of DFUs and accelerate the wound healing process, thereby providing a safer, more economical, and more effective option for the clinical management of DFUs.

## Material and Methods

### Participants in moderate-intensity aerobic exercise

This study enrolled five healthy adult female volunteers, aged 22–30 years, with a body mass index (BMI) of 18–25 kg/m^2^ and no history of systemic diseases. In the month prior to enrollment, none of the participants had taken any medication known to affect sweating or inflammatory responses. Following a standardized protocol, all participants underwent a thorough pre-examination screening, which included collection of basic demographic information (Table S1), completion of a Physical Activity Readiness Questionnaire (PAR-Q), and baseline medical checks such as blood pressure measurement and routine blood tests. This screening ensured that every participant could safely complete the moderate-intensity aerobic exercise session for sweat collection.

The study purpose, procedures, sample-collection methods, potential risks and benefits, and privacy safeguards were explained to all participants in clear language. Each participant provided written informed consent. Throughout the study, biological sample collection and data handling strictly followed ethical requirements. Participants were free to withdraw at any time without any negative consequences.

All study procedures were approved by the Ethics Committee of Wuhan Sports University (Approval No. 2025089) prior to study initiation.

### Exercise protocol

As none of the participants had a history of regular exercise training, this study estimated each individual’s maximum heart rate (HRmax) using the Tanaka formula :HRmax = 208 − (0.7 × age)^26^.

Based on this, the target exercise intensity was set at 60%–80% of the individual’s HR_max to ensure a moderate-intensity range^27^. Based on participants’ ages, the resulting target heart rate range was approximately 116–138 beats per minute.

To minimize the influence of potential confounding factors on sweat secretion and metabolic state, all experimental sessions were scheduled outside the participants’ menstrual periods. For 24 hours prior to the exercise intervention, participants were required to strictly adhere to a standardized pre-experimental protocol. This included abstaining from alcohol, coffee, strong tea, and other caffeinated beverages, and avoiding high-sugar, high-fat, spicy, or stimulant-rich foods, and fasting for at least 2 h before the experiment. These measures aimed to reduce variability in metabolic background caused by dietary differences and to prevent potential interference with subsequent biofluid-related experiments.

The specific exercise protocol was as follows: Participants first performed a 5-minute progressive warm-up on a treadmill, followed by 30 minutes of continuous running while maintaining their individual target heart rate range^27^. Throughout the entire session, heart rate was monitored continuously and in real time using a Polar H10 heart rate sensor to ensure the exercise intensity consistently met the preset requirements.

### Sweat collection and sample processing

Sweat was collected using an absorbent filter paper patch method. Prior to sampling, the skin on the participant’s back was cleansed with alcohol and deionized water and dried with sterile gauze. Two pieces of absorbent filter paper (Whatman™ quantitative filter paper, Grade 1825-025, Cytiva, UK), each 25 mm in diameter, were placed on the skin and covered with a waterproof transparent medical adhesive dressing (3M™ Tegaderm™ transparent film dressing, model 1624W, 3M Health Care, USA) to secure them in place. Throughout the procedure, the researchers wore disposable gloves and handled the filter paper only with sterile forceps to minimize exogenous contamination.

Sweat collection lasted approximately 25 minutes to obtain sufficient volume for subsequent analysis. During this period, the patch was visually inspected at regular intervals; if the filter paper appeared close to saturation, it was removed early to prevent sweat overflow or sample loss.

After collection, the filter paper was immediately transferred to a pre-cooled nuclease-free centrifuge tubes. For elution, 2 mL of normal saline was added per filter paper piece, and the tubes were agitated on an orbital shaker at 4 °C and 200 rpm for 2 h. The eluate was then centrifuged at 4 °C and 4000 rpm for 5 min. The supernatant was collected and passed through a 0.22 μm syringe filter to obtain a clear sweat sample. Samples were stored in an ultra-low-temperature freezer (−80 °C) until further analysis.

During storage, water evaporation may lead to overestimation of solute concentration in sweat samples^28^. To reduce this bias, following previous recommendations^28,29^, samples were sealed with impermeable Parafilm® M film to improve sample stability and the reliability of analytical results.

### Cell culture and treatment

In order to obtain the transcriptome induced by the exercise-indued sweat, we select the A549 cell line as the tool cell and treated A549 with the sweat and the control. The A549 cell line was routinely cultured in high-glucose DMEM supplemented with 10% fetal bovine serum and 1% penicillin-streptomycin, and maintained at 37 °C in a humidified atmosphere with 5% CO_2_. Cells in the logarithmic growth phase were seeded into 6-well plates. When reaching 70–80% confluency, the medium was replaced with fresh medium containing either post-exercise sweat (HE) or sweat-saline control (Hcon) at an equal mass, followed by incubation for 24 h. Three independent biological replicates were performed for each experimental group.

### RNA sequencing and transcriptomic Analysis

Total RNA was isolated from A549 cells using Trizol reagent (Invitrogen). RNA quality and integrity were assessed with an Agilent 2100 BioAnalyzer and a Qubit Fluorometer; only samples with an RNA integrity number (RIN) > 7.0 and a 28S:18S ratio > 1.8 were used for downstream processing.

RNA-seq libraries were constructed by CapitalBio Technology (Beijing, China) using the NEB Next Ultra RNA Library Prep Kit for Illumina. Briefly, poly(A)+ mRNA was enriched from 1 µg of total RNA, fragmented, and converted into double-stranded cDNA. After end repair, A-tailing, and adapter ligation, libraries were amplified by PCR. Final library quality and concentration were validated using an Agilent 2100 Bioanalyzer and the KAPA Library Quantification kit. Libraries were then subjected to paired-end 150 bp sequencing on an Illumina NovaSeq platform.

For data analysis, raw read quality was assessed with FastQC (v0.11.5) and adapters/low-quality bases were trimmed using NGSQC (v2.3.3). Cleaned reads were aligned to the human reference genome (hg38) using HISAT2 (v2.1.0). Transcript assembly and gene-level expression quantification were performed with StringTie (v1.3.3b). Differential expression analysis was conducted using DESeq2 (v1.28.0).

We selected the genes with a fold change (FC) ≥ 1.50 or ≤ 0.67 as the gene signature induced by sweat. Finally, we used an in-house computer program to predict the potential protective roles of exercise-induced sweat based on the reverse transcriptomics approach.

### In vivo wound healing study in diabetic foot ulcer animal model

#### Experimental animals

Male 7-week-old db/db mice (n=15) were purchased from the Model Animal Research Center of Changzhou Cavens (Jiangsu, China). All procedures were carried out according to the rules of the Committee on Animal Care and Use published by the Institute of Chinese Materia Medica, China Academy of Chinese Medical Sciences. The mice were anesthetized with tribromoethanol (20 ml/kg, intraperitoneal injection, i.p.) and then wound molding treatment after back hair shaving. A circular 6 mm full-thickness cutaneous wounds on the back of mice were made using a sterile 6-mm biopsy punch, and 0.5 mm-Thick silicone donut-shaped splints were fixed around the wounds and positioned with 4-0 nylon sutures. The mice were randomly allocated to different groups to receive either physiological saline, post-exercise sweat (original solution, 100μl/day, topical), or rhEGF (4000 IU/10*10cm2, topical) until study end. The wounds were photographed at 0, 3, 7, 10, and 14 days after surgical wounds were made. The wound closure photos were analyzed using ImageJ by calculating the percentage of the wound area from the original wound area. After 14 days, skin tissues were collected which were paraffin-embedded for histology.

#### Histology, immunohistochemistry and Immunofluorescence Staining

The paraffin tissue section slides with thickness of 5 μm were used for histology and immunofluorescence staining. After deparaffinized, the slides were stained with hematoxylin and eosin (HE) and Masson’s trichrome stain and following their instructions, respectively. IHC staining using primary antibodies against ki67 (1:200, abcam). For immunofluorescence (IF) staining, the sections were blocked by 5% (m/v) BSA, after they were permeabilized with 0.5% Triton X-100. Primary antibody was loaded and kept at 4°C overnight. Then Alexa Fluor 488- and Alexa Fluor 647-conjugated secondary antibodies were loaded at room temperature for 1 h. The nuclei were stained with 4′,6-diamidino-2-phenylindole (DAPI). The images were visualized using microscope.

## Results

### Predicting exercise-induced sweat to treat diabetic foot ulcers

As shown in Figure 1, we proposed a novel sports medicine paradigm, that is, exploring the potential protective roles of exercise-induced sweat. We first collected the sweat sample induced by moderate-intensity aerobic exercise from five young female participants. We then used the sweat to treat the A549 cell line and subsequently identified the transcriptomic gene signature induced by sweat using RNA-sequencing. Finally, the protective role of sweat was predicted by an in-house computer program based on reverse transcriptomics approach.

**Figure 1.**
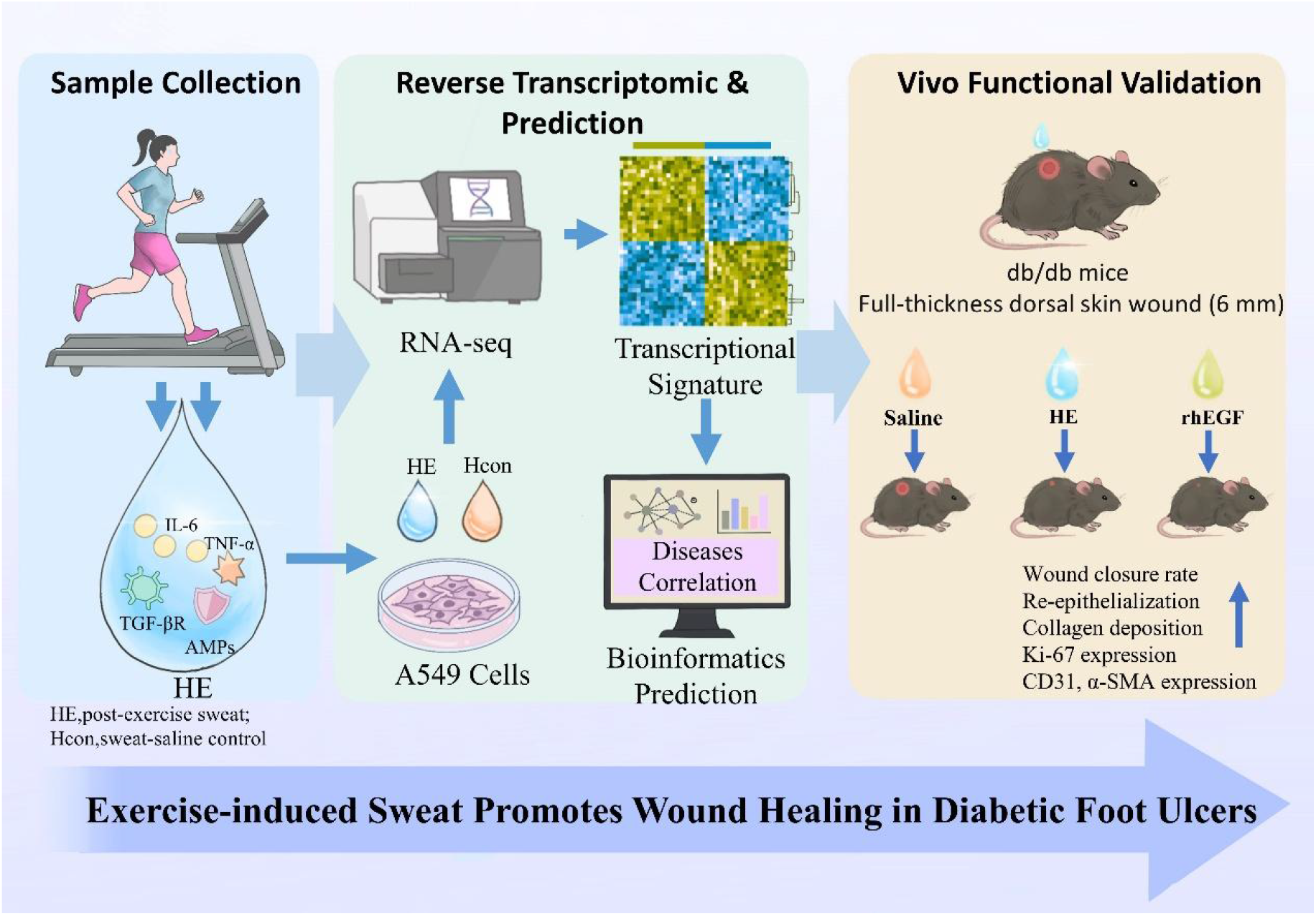
Overview of the study design and experimental workflow.

As a result, DFUs is one of the most significantly (top 3) diseases predicted to be treated by exercise-induced sweat (OR = 0.57, p-value = 7.2e-4, Fisher’s Exact test). We identified 188 down-regulated genes in DFUs are upregulated by exercise-induced sweat, whereas 199 up-regulated genes in DFUs are downregulated by exercise-induced sweat (**Figure 2a**). Moreover, the WEAT tool analysis revealed that these genes are mostly enriched in biological processes (BP) such as positive regulation of glucose catabolic process to lactate via pyruvate (p-value = 0), water transport (p-value = 1.43e-4), and neutrophil aggregation (p-value = 2.00e-4), which plays critical roles in inflammatory response, pathogen clearance, and would healing (**Figure 2b**). The above results together suggest that the sweat induced by moderate-intensity aerobic exercise may have protective roles in preventing DFUs.

**Figure 2.**
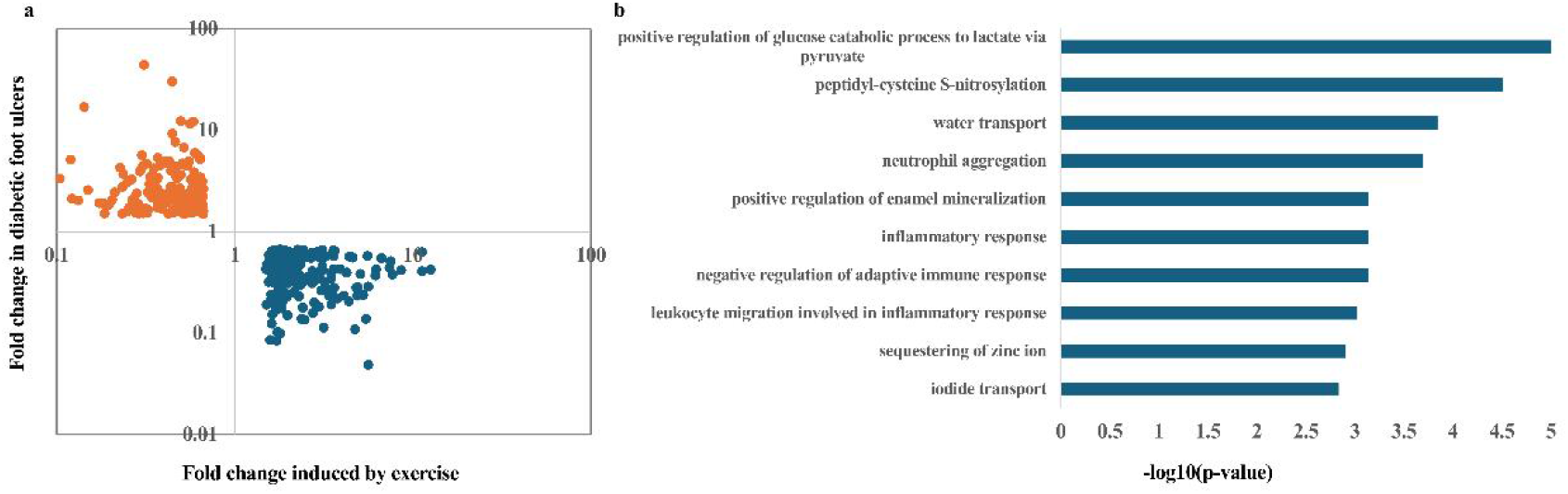
Scatter plots (a) and the top 10 mostly enriched biological processes (b) of the genes dysregulated in diabetic foot ulcers and reversed by exercise-induced sweat.

### Animal experiments confirming exercise-induced sweat can promote would healing of diabetic foot ulcers

To confirm the therapeutic potential of moderate-intensity aerobic exercise induced sweat in DFUs, we generated full-thickness wound models on the dorsal skin of diabetic mice and treated wounds with saline, sweat, or recombinant bovine basic fibroblast growth factor gel (rhEGF) (**Figures 3a**). As expected, mice treated with sweat exhibited a lower percentage of unclosed wounds on days 3, 7, 10, and 14 post-wounding, with significantly lower values relative to the control group on days 10 and 14. By day 14, wounds in the sweat-treated group were nearly completely healed, with a wound closure rate of 97%, whereas large wound areas remained in the control group (**Figures 3b-c**). These results indicate that sweat treatment markedly accelerates diabetic wound healing. Of note, the percentage of unclosed wounds in the rhEGF-treated group was lower than that in the control group and remained comparable to that in the sweat-treated group throughout the observation period. This finding suggests that sweat treatment exerts therapeutic effects similar to those of rhEGF.

**Figure 3.**
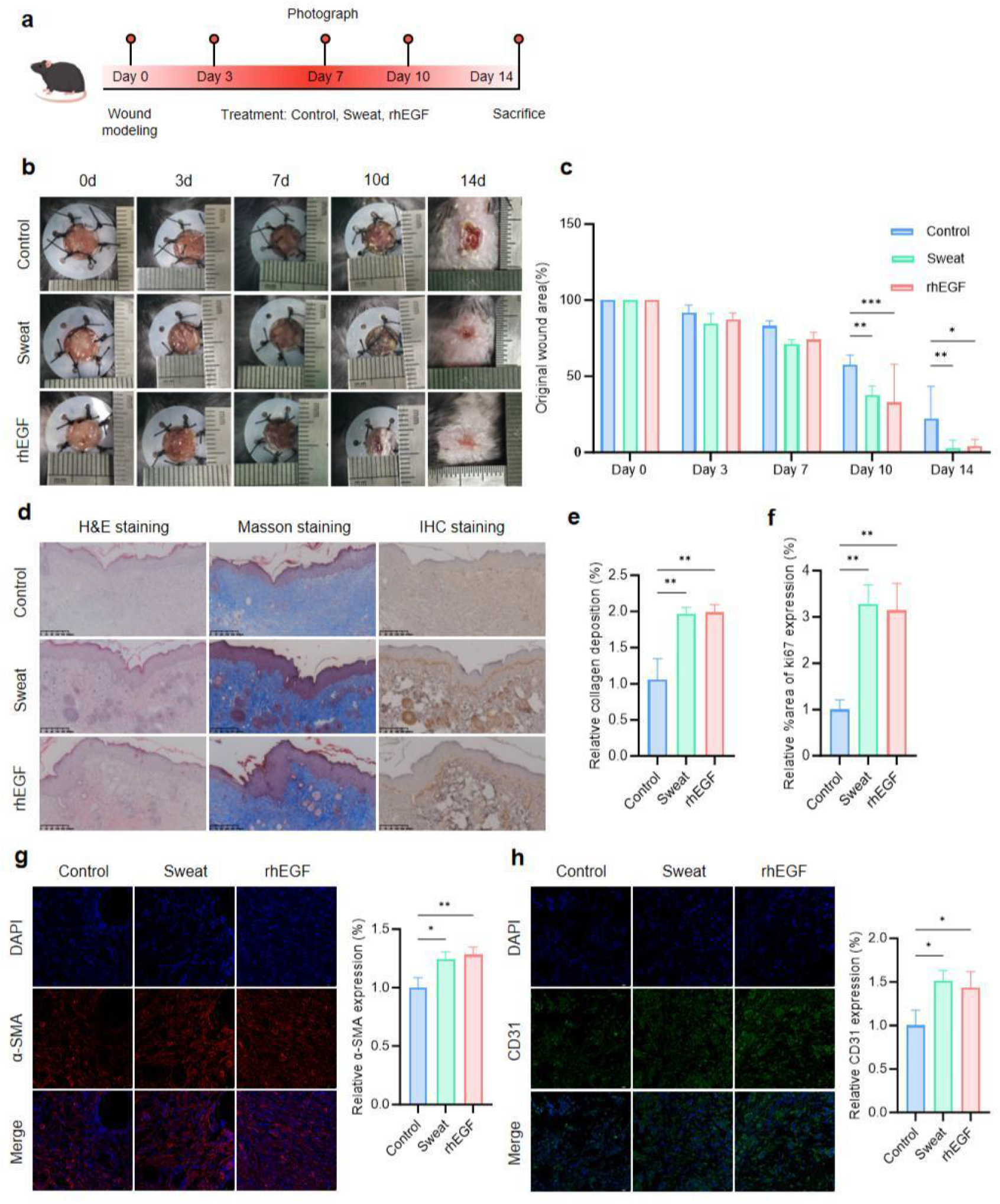
Athlete sweat improves wound healing in db/db mice. Full-thickness wounds were created on the dorsal skin of db/db mice, and normal saline, athlete sweat, or recombinant bovine basic fibroblast growth factor gel was applied to the wound edges as previously described. (a) Schematic illustration of the in vivo mouse experiment. (b) Representative wound images (n=5). (c) Percentage of initial wound area (n=5). (d) Representative images of H&E, Masson’s trichrome staining, and IHC of dorsal skin sections; scale bar = 200 μm. (e) Quantification of collagen deposition by Masson’s trichrome staining. (f) Quantification of Ki67 expression from IHC images. (g) Representative α-SMA immunofluorescence images and quantification of dorsal skin sections; scale bar = 50 cm. (h) Representative CD31 immunofluorescence images and quantification of dorsal skin sections; scale bar = 50 μm. (n = 3). Data are presented as mean ± SD. Statistical analyses were performed using one-way ANOVA and two-way ANOVA. \**P* < 0.05, \*\**P* < 0.01, indicating statistical significance.

Subsequent histological analysis revealed poor wound healing in the control group, with incomplete epithelial coverage of the wound and marked inflammatory cell infiltration. In contrast, wounds in the sweat-treated and rhEGF-treated groups displayed complete re-epithelialization and reduced inflammatory cell infiltration. Furthermore, collagen deposition and Ki67 expression were higher in the sweat-treated group than in the negative and positive controls, suggesting that sweat treatment may promote wound healing in diabetic mice (**Figure 3d–1f**). CD31 and α-SMA serve as two vascular markers to evaluate angiogenesis in the diabetic wound bed. Immunofluorescence results showed that only a few vessels were observed in wound tissues of the control group, whereas treatment with sweat and rhEGF improved vascular network formation. Of note, vascular density was high and comparable between the sweat-treated group and the rhEGF-treated group (**Figure 1g-h**). Collectively, these data indicate that sweat may significantly accelerate diabetic wound healing by promoting wound re-epithelialization, collagen deposition, and angiogenesis.

## Discussion

Although a broad spectrum of therapeutic agents is currently available for diabetes management, clinical care of DFUs still lacks safe, cost-effective, and highly potent treatment strategies. Therefore, beyond glycemic control, identifying agents that effectively attenuate DFUs progression and accelerate wound healing is crucial to further improving the quality of life of diabetic patients and alleviating the national healthcare burden. The present study is the first to demonstrate that, in a murine model of diabetic wounds, sweat treatment significantly enhances wound healing, re-epithelialization, and collagen deposition compared with control treatment. Concomitantly, the expression levels of the proliferation marker Ki67 and the angiogenesis markers CD31 and α-SMA were markedly upregulated. These findings suggest that sweat not only contributes to cutaneous innate immunity but also targets the pathophysiological deficits underlying DFUs, thereby exerting robust pro-healing effects. This work provides a novel theoretical framework for developing sweat as a natural therapeutic agent for diabetic complications.

This study has several limitations that warrant consideration. First, sweat treatment was only evaluated in an animal model of diabetic wounds, with analyses focused on phenotypic outcomes rather than the key bioactive factors and molecular mechanisms mediating the pro-healing effects. Future investigations will therefore focus on deciphering these underlying mechanisms. Second, the sweat used in this study was collected exclusively from females with a relatively small sample size, and male donors were not included. Given that sweat composition is known to be influenced by sex, age, physical activity level, and physiological status ^17,30,31^, the generalizability of our findings may be limited. Future studies should expand the donor pool to include males, individuals of different age groups to validate the consistency of sweat’s pro-healing efficacy. Third, this study was restricted to an in vivo murine model, and the translational potential of sweat therapy for human DFUs remains to be confirmed. Follow-up studies should include in vitro experiments to verify the direct effects of sweat on cell proliferation, migration, and angiogenesis. Additionally, this study only explored sweat induced by moderate-intensity aerobic exercise, the therapeutic effects remain unclear for other types of exercises. Finally, well-designed clinical trials are warranted to assess the safety, efficacy, and optimal administration route of sweat in patients with DFUs, while accounting for variables such as wound type and patient comorbidities.

In summary, the present study provides preliminary evidence that exercise-induced sweat treatment promotes diabetic wound healing in mice. Given its natural origin, inherent bioactivity, low cost, and high biocompatibility, sweat emerges as a promising candidate for the development of novel, safe, and cost-effective therapeutics for DFUs. Addressing the current limitations through in-depth mechanistic dissection, expansion of the donor cohort, and translational research will be critical to advancing this innovative strategy toward clinical application—ultimately offering a new therapeutic option to improve the clinical management of DFUs.

## Acknowledgements

This study was supported by the Noncommunicable Chronic Diseases-National Science and Technology Major Project (2024ZD0531201) and Natural Science Foundation of China [82427801, 32301239] and the Scientific and Technological Research Project of Xinjiang Production and Construction Corps (2023AB057) and the Natural Science Foundation of Hubei Province (2025AFD622).

## Completing interests

The authors declare no competing interests.

## Author Contributions

QC proposed the project. HY, QF, and MS determined the exercise protocol. JZ and CW designed the animal experiments. MZ and YT performed the animal experiments. HY, JC, and TL performed experiments of RNA-sequencing and reverse transcriptomics. FQ, ML and BY took part in data analysis. MZ, JZ, HY and QC wrote the draft manuscript. CW, JZ, and QC supervised the study.

